# Spironolactone but not Eplerenone Exacerbates Cisplatin Nephrotoxicity

**DOI:** 10.1101/790998

**Authors:** Gabriel R. Estrela, Benjamin Bonnard, Jonatan Barrera-Chimal, Frédéric Jaisser

## Abstract

Cisplatin is a highly successful chemotherapeutic agent used for the treatment of solid tumors. However, nephrotoxicity is a limiting factor that occurs in 30% of patients under treatment. Many mechanisms are involved in cisplatin-induced nephrotoxicity, such as epithelial and endothelial injury, inflammation, oxidative stress, and renal vasoconstriction. The mineralocorticoid receptor (MR) has an important role in inflammation and vascular function. MR blockage and ablation have been shown to be effective in preventing renal ischemia-reperfusion injury and cyclosporine A-induced nephrotoxicity. We investigated whether MR antagonism with spironolactone or eplerenone could prevent cisplatin-induced nephrotoxicity. Here, we show that spironolactone treatment exacerbates nephrotoxicity in mice treated with acute and long-term cisplatin regimes. Moreover, spironolactone potentiated the toxicity induced by cisplatin treatment in a cell viability assay in human embryonic kidney cells. In contrast, eplerenone neither prevented nor increased cisplatin toxicity in mice or cultured cells. Thus, our studies support recent findings showing that spironolactone potentiates cisplatin-induced cytotoxicity, independently of mineralocorticoid receptor inhibition.

## Introduction

Cisplatin (CIS) is an antineoplastic drug used in chemotherapy with a high rate of success to treat solid tumors. However, its use is limited by its side effects, especially nephrotoxicity, which affects 30% of patients treated with CIS (1, 2). CIS-induced nephrotoxicity is due to several mechanisms, such as direct epithelial and endothelial injury, renal vasoconstriction, inflammation, oxidative stress, hypoxia, and apoptosis (2–4). Mineralocorticoid receptor (MR) antagonism is widely used for the treatment of cardiovascular complications of heart failure (5–7). Our group demonstrated that MR antagonism protects against acute kidney injury (AKI) induced by Cyclosporin A (8) and ischemia-reperfusion, two conditions characterized by sustained vasoconstriction, resulting in epithelial and endothelial injury (9–11). MR antagonism also attenuates the AKI-to-chronic kidney disease transition in a bilateral renal ischemia-reperfusion model (12–14). These observations in rodent models have been extended to the large white pig ischemia-reperfusion model (11, 13).

Our main objective was to analyze whether pharmacological MR antagonism can protect kidneys against nephrotoxicity induced by CIS.

## Materials and Methods

### Experimental protocols

All procedures used and described for animal experiments were carried out in accordance with the National and European guidelines for animal experimental research and were approved by the Ethical Committee for Animal Experiments of the Sorbonne University (#7594), France. The animals were housed in controlled-climate conditions with a 12-hour light-dark cycle and provided free access to food and water. Eight-week-old male C57Bl/6 mice (Janvier Laboratories, France) were used for pharmacological experiments with spironolactone (Sigma-Aldrich), eplerenone (Inspra, Pfizer), and CIS (Sigma-Aldrich). Spironolactone (20 mg/kg) was administered by oral gavage (vehicle: methylcellulose) in a single dose, 48, 24, and 1h before each CIS injection. Eplerenone (200 mg/kg/day) was administered by oral gavage in two doses of 100 mg/kg, 48, 24, and 1 h before CIS injection. The doses of spironolactone and eplerenone were chosen according to our past experience and based on previously published studies in mice (15–23). Acute kidney injury was induced with one single dose of CIS (20 mg/kg) i.p and mice were euthanized three days later. Long-term CIS treatment consisted of repeated dosing with one dose of CIS (7 mg/kg) i.p per week for four weeks. Mice were euthanized three days after the last injection.

### Renal Parameters analysis

Blood sample was taken via cardiac punction and plasma urea and creatinine concentrations were determined with an automatic analyzer (Konelab 20i, Thermoscientific). Urine was collected in metabolic cages over 24 hours after 3 days period of habituation and protein concentration were determined with Konelab 20i.

### RNA extraction and real-time PCR

Total RNA was extracted from the kidneys with TRIZOL reagent (Life Technologies, Carlsbad, CA) according to the manufacturer’s instructions. Reverse transcription was performed with 1 μg RNA and the Superscript II Reverse Transcriptase kit (Life Technologies). Transcript levels were analyzed by real-time polymerase chain reaction (RT-PCR) (fluorescence detection of SYBR green) in an iCycler iQ apparatus (Bio-Rad, Hercules, CA), with normalization against 18S as an endogenous control. The primer sequences for the analyzed genes are listed in Supplementary Table 1.

### Histological analysis

The kidneys were fixed in Bouin’s fixative solution and then dehydrated and embedded in paraffin. Sections (4 μm) were cut and stained with Hematoxylin Eosin and Sirius Red. At least six subcortical fields were visualized and analyzed for each mouse using a Leica DM4000 microscope at a magnification of 200X. The acute tubular necrosis score was determined based on the percentage of tubules showing luminal casts, cell detachment, or dilation and assigned according to the following scale: 0 = 0 to 5%, 1 = 6 to 25%, 2 = 26 to 50%, 3 = 51 to 75%, and 4 > 75%. Histology analysis was performed blind to experimental groups to assess tubule-interstitial fibrosis based on the Sirius Red–positive area and assigned according to the following scale: 1 ≤ 25%, 2 = 26 to 50%, 3 = 51 to 75%, and 4 > 75%.

### Cell treatments

Human embryonic kidney (HEK) 293 cells were cultured in Dubelcco’s modified Eagle’s medium (DMEM; 31966, *ThermoFisher*), 10% fetal bovine serum (FBS; S1810, *Dutcher*), and 1% Penicillin/Streptomycin (P/S; 151140122, *ThermoFisher*) at 37°C in 5% CO_2_ and used between passages 7 and 9. At 90% confluence, the cells were exposed to 0.05% trypsin-EDTA (25300054, *ThermoFisher*) and seeded in 96-well plates at 2,000 cells per well. After 48 h, cells were starved with DMEM containing 1% FBS and 1% P/S for 24 h. the cells were treated with various concentrations of cis-Diammineplatinum(II) dichloride (25, 40, or 60 μM; P4392, *Sigma*) diluted in 0.9% NaCl solution, with or without spironolactone (10 μM; S3378, *Sigma*) (24) diluted in 1% ethanol or eplerenone (100 or 200μM; 12929536002, *AstraZeneca*) (25) diluted in 2% DMSO. Cells were treated for 24 h.

### Cell viability assay

After treatment, cell viability was measured with a cell proliferation kit (MTT, 1146500700, *Sigma*) according to the manufacturer’s instructions. Briefly, MTT labeling reagent was added to the cells in culture media, the cells incubated for 4 h at 37°C, solubilization solution added, and the cells further incubated overnight at 37°C. The absorbance values were obtained at 550 nm using an Infinite 200 Pro (*Tecan*).

### Statistics

The results are expressed as the mean +/− SEM. The significance of between-group differences was assessed by one-way analysis of variance (ANOVA) with the Tukey correction for multiple comparisons. A Student t-test was performed to compare two groups. GraphPad (La Jolla, CA) Prism 6 software was used for statistical analyses and graphical representations. p values < 0.05 were considered to be statistically significant.

## Results

### Spironolactone but not Eplerenone exacerbates renal dysfunction and renal injury after single-dose administration of cisplatin

Plasma creatinine and plasma urea levels were higher in the CIS group than vehicle (VEH) group and further exacerbated in the CIS+SPIRO group (Figure 1A-B). The mRNA expression of NGAL, a renal injury marker, and TNF-α, an inflammatory marker, was also higher in the CIS group than VEH group and further elevated in the CIS+SPIRO group (Figure 1C-D). Tubular injury, assessed by HE staining, was higher in the CIS group than VEH group and further exacerbated in the CIS+SPIRO group (Figure 1E-H). In contrast to spironolactone, eplerenone neither exacerbated nor protected against the deleterious effects of CIS (Figure 2A-H).

**Figure 1.**
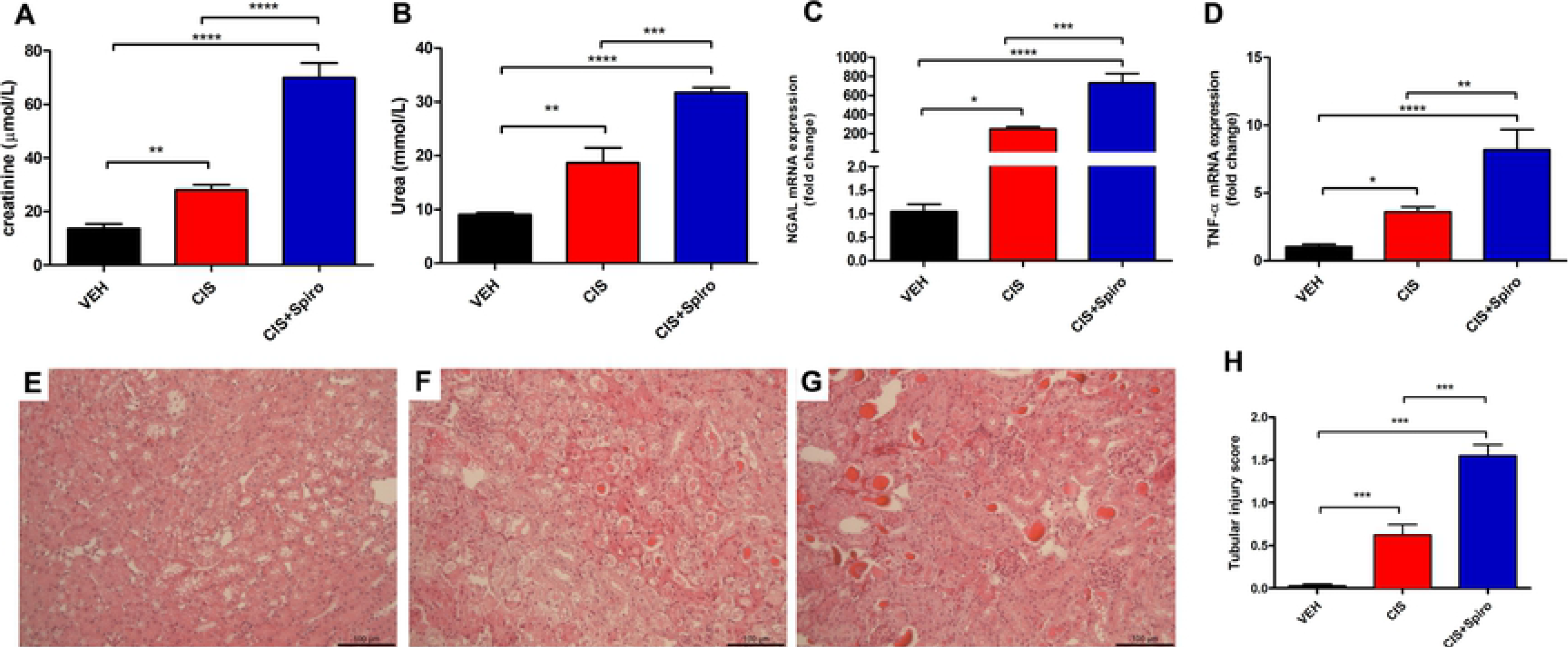
Spironolactone exacerbates acute cisplatin-induced nephrotoxicity. Renal function was determined by quantifying the plasma levels of (**A**) creatinine and (**B**) urea. Tubular injury was determined by (**C**) NGAL and (**D**) TNF-α mRNA expression. Representative H&E staining is shown for mice treated with (**E**) vehicle (VEH), (**F**) CIS alone (CIS), or (**G**) CIS plus spironolactone (CIS+SPIRO). (**H**) Acute tubular necrosis score. n = 6 to 7 per group. One-way analysis of variance was performed. Bar = 100 μm. *P < 0.05, **P < 0.01, ***P < 0.001, ****P < 0.0001.

**Figure 2.**
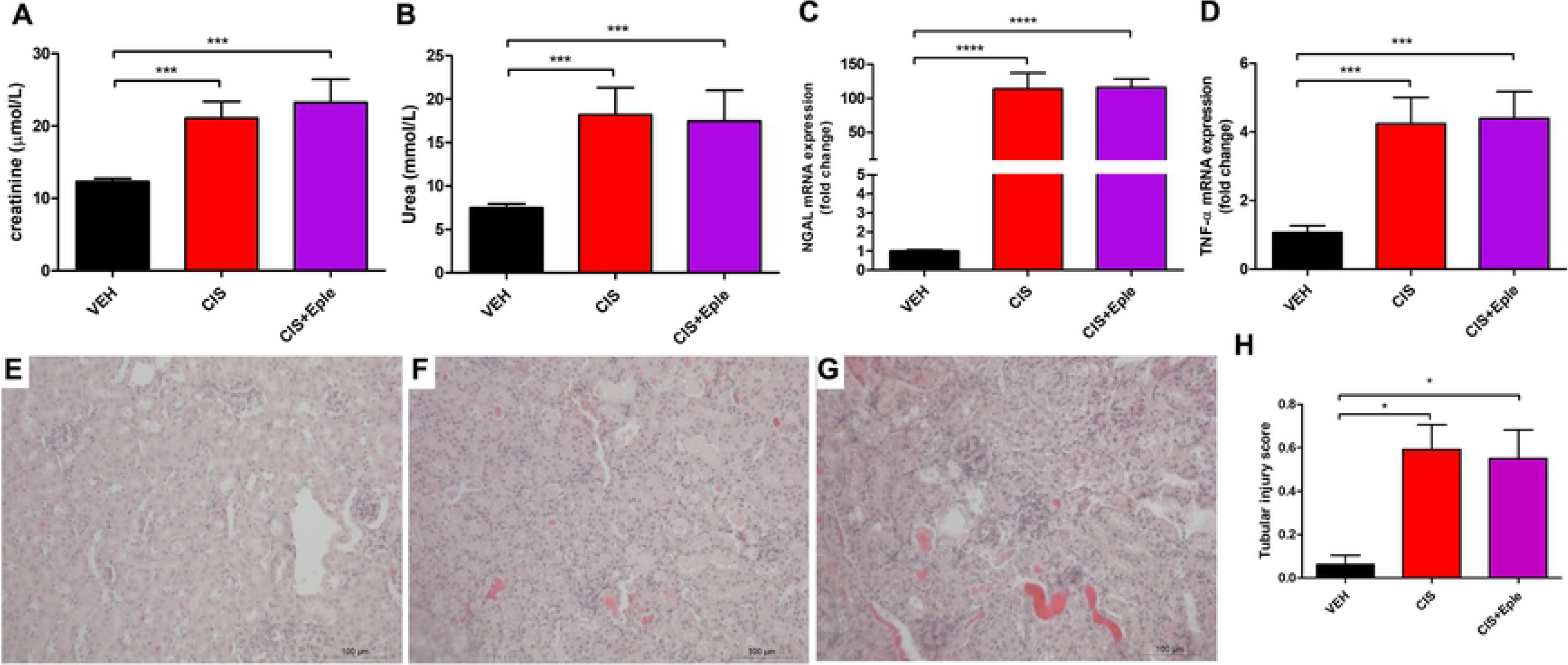
Eplerenone does not alter acute cisplatin-induced nephrotoxicity. Renal function was determined by quantifying the plasma levels of (**A**) creatinine and (**B**) urea. Tubular injury was shown by (**C**) NGAL and (**D**) TNF-α mRNA expression. Representative H&E staining is shown for mice treated with (**E**) vehicle (VEH), (**F**) CIS alone (CIS), or(**G**) CIS plus eplerenone (CIS+EPLE). (**H**) Acute tubular necrosis score from H&E-stained sections. n = 6 to 7 per group. One-way analysis of variance was performed. Bar = 100 μm. *P < 0.05, **P < 0.01, ***P < 0.001, ****P < 0.0001.

### Spironolactone exacerbates nephrotoxicity after long-term cisplatin treatment

In clinical practice, patients receive CIS at low doses repeatedly and for long periods. We thus developed a new model of repeated low-dose CIS administration for four weeks. Long-term CIS treatment induced renal dysfunction, with increased polyuria and proteinuria (Figure 3A-C). Combined treatment with CIS and SPIRO significantly increased plasma creatinine levels relative to those of the VEH group (Figure 3A-C). The renal expression of TGF-β and Col4 was unchanged (Figure 3D and E), whereas the mRNA levels of NGAL and TNF-α were induced by CIS (Figure 3 F-G). Long-term CIS induced moderate interstitial fibrosis in the absence of clear tubular injury (Figure 3K-0). Spironolactone co-treatment exacerbated proteinuria (whereas the level of increased plasma creatinine and polyuria was unchanged) (Figure 3 A-C), as well as the renal expression of TGF-β, Col4, NGAL, and TNF-α (Figure 3 D-G). Importantly, spironolactone potentiated interstitial fibrosis and led to a high level of tubular injury relative to that of the CIS or VEH groups (Figure 3 K-O). Thus, spironolactone co-treatment clearly potentiated long-term CIS-induced nephrotoxicity.

**Figure 3.**
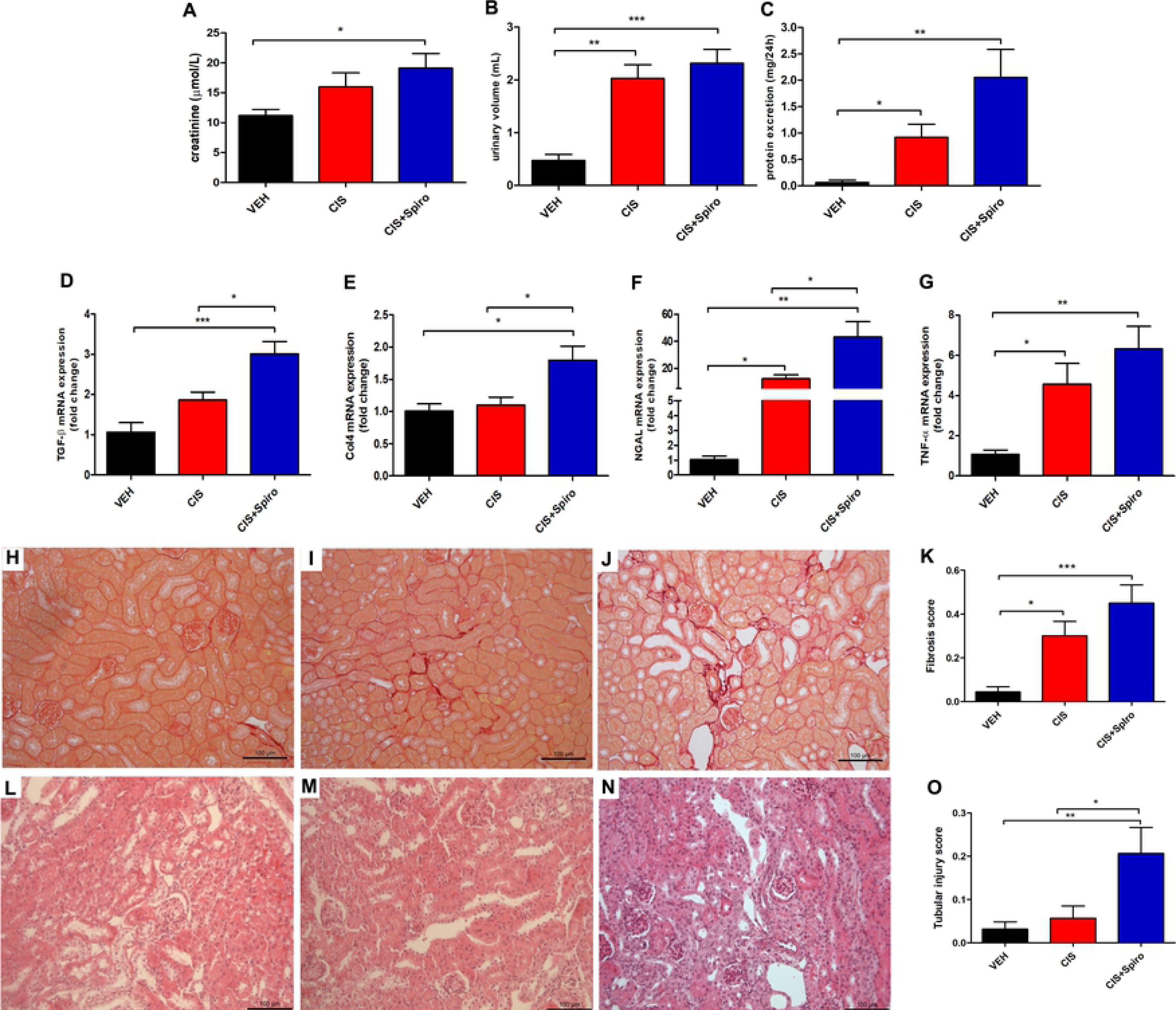
Spironolactone exacerbates long-term cisplatin-induced nephrotoxicity. Renal function was determined by quantifying the plasma levels of (**A**) creatinine (**B**) urinary volume, and (**C**) protein excretion. We determined the kidney mRNA levels of (**D**) transforming growth factor and (**E**) collagen IV by real-time PCR as markers of kidney fibrosis and (**F**) NGAL and (**G**) TNF-α mRNA levels as markers of tubular injury. Representative Sirius Red–stained images for the (**H**) vehicle, (**I**) CIS treated (CIS), and (**J**) CIS plus spironolactone treated (CIS+SPIRO) groups. (**K**) Fibrosis score estimated from Sirius Red staining. Representative H&E staining is shown for mice treated with (**L**) vehicle (VEH), (**M**) CIS alone (CIS) or (**N**) CIS plus spironolactone (CIS+SPIRO). (**O**) Acute tubular necrosis score from H&E-stained sections. n = 6 to 7 per group One-way analysis of variance was performed. Bar = 100 μm. *P < 0.05, **P < 0.01, ***P < 0.001, ****P < 0.0001.

### Spironolactone but not eplerenone decreases HEK-293 cell viability after cisplatin exposure

We next assessed cell viability by the MTT assay. HEK cells were incubated with spironolactone or eplerenone for 24 h and then exposed to CIS for 24 h. CIS decreased cell viability for all doses tested (25/40/60 μM) (Figure 4A). The diluent of CIS, spironolactone or eplerenone (NaCl, EthOH or DMSO, respectively) has no effect on viability (Figure 4A-B). The addition of spironolactone decreased cell viability for all tested concentration of CIS (Figure 4A). Importantly, eplerenone at 100 orr 200 microM did not affect the cell toxicity of CIS, in contrast to spironolactone (Figure 4B).

**Figure 4.**
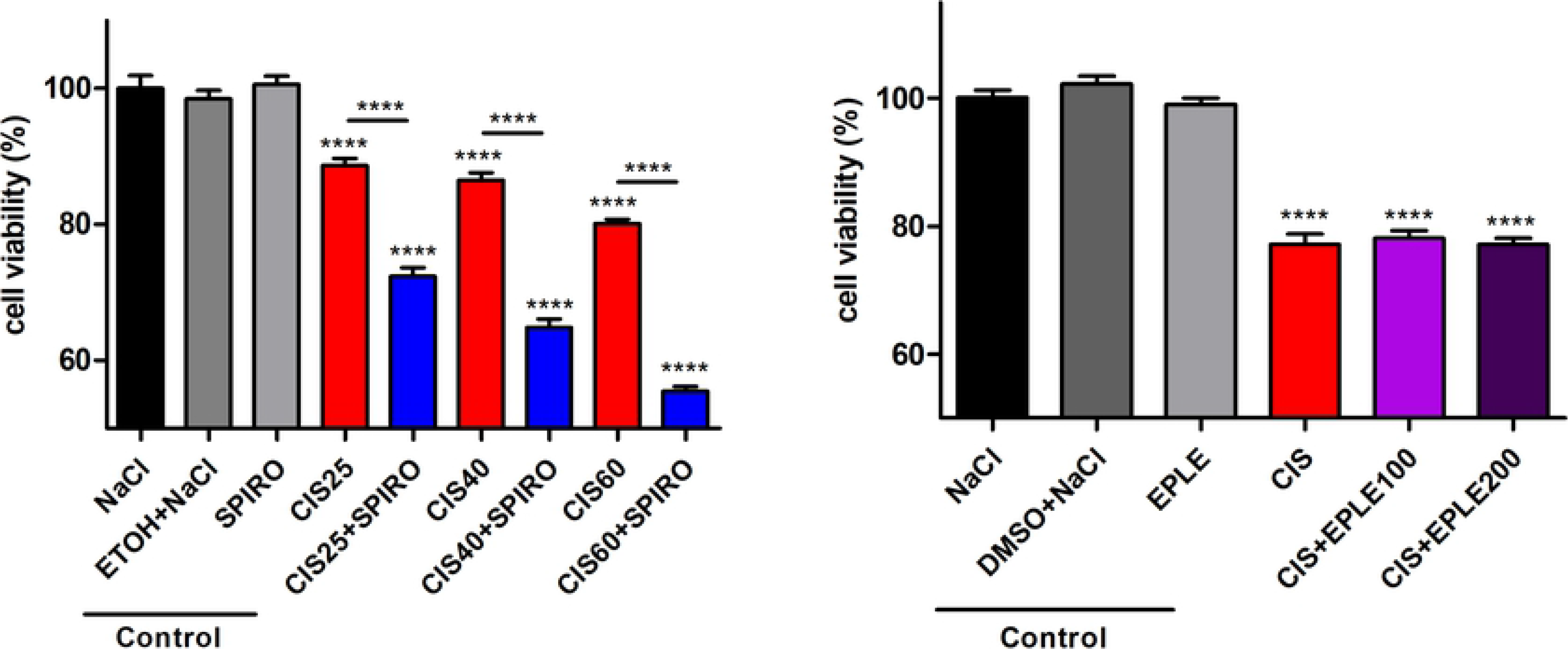
Spironolactone but not eplerenone decreases HEK 293 cell viability. (**A**) HEK 293 cells were treated with 25, 40, or 60 μM CIS alone or together with 10 μM spironolactone for the cell viability assay, (**B**) HEK 293 cells treated with 40 μM CIS alone or together with 100 or 200 μM eplerenone. One-way analysis of variance was performed. ****P < 0.0001 vs control groups <u>****</u>P < 0.0001 CIS vs CIS+SPIRO.

## Discussion

Approximately one-third of patients treated with CIS develop nephrotoxicity (1, 2). CIS enters renal cells by a facilitated mechanism or a passive route and activates pathways that promote cell death (1, 2, 26). In addition, CIS induces TNF-α production by tubular cells, triggering an inflammatory response which contributes to tubular cell injury and death (27, 28). The renal vasculature is also targeted, resulting in ischemic tubular cell death (2). Part of the nephrotoxicity of CIS is due to vascular damage and vasoconstriction, caused mainly by endothelial dysfunction and abnormal vascular autoregulation (29). CIS can also be directly toxic to the vascular endothelium (29). Endothelial injury results in overexpression of endothelial cell-adhesion molecules and the binding of leukocytes (30). Vascular congestion and leukocyte infiltration result, leading to the injury and swelling of endothelial cells. Moreover, such obstruction decreases renal blood flow, contributing to the local production of vasoconstrictors and affecting the glomerular filtration rate (29, 31).

Our group previously demonstrated that MR antagonism attenuates kidney injury in various experimental models. Pharmacological MR antagonism by all current classes of MR antagonists (steroidal, such as spironolactone or eplerenone, or non-steroidal, such as BR-4628 and finerenone) protects against acute kidney injury induced by ischemia-reperfusion in mice, rats, and large white pigs (10, 11, 14, 32) as well as cyclosporine-induced nephrotoxicity (8, 10–13). We hypothesized that a similar benefit of MR antagonism might be observed for acute CIS-induced nephrotoxicity, as several common underlying mechanisms have been proposed, including altered hemodynamics, oxidative stress, and an inflammatory response. Indeed, pharmacological MR antagonism during renal ischemia-reperfusion has highlighted the role of MR-mediated oxidative stress in the acute phase of ischemia (10, 11), which leads to decreased renal perfusion and increased renal resistance. Similar observations were made in an acute model of cyclosporin-induced nephrotoxicity, in which MR deletion in smooth muscle cells improved renal perfusion, thus attenuating vascular resistance and modulating L-type Ca^2+^ channels (8). Importantly, blunting MR activation in the early phase of IRI (by short-term treatment with MRAs) in rodents or large white pigs has long-term benefits, even in the absence of continued MRA administration, preventing the development of chronic kidney disease with impaired renal function, proteinuria, and kidney fibrosis (13). In the present study, we show that these benefits do not extend to CIS-induced nephrotoxicity, since both acute and chronic CIS-induced nephrotoxicity were not blunted by MRAs given just before and at the time of CIS administration, when the acute phase of nephrotoxicity maybe maximal. Indeed, we observed the opposite effect with spironolactone, as it aggravated CIS-induced nephrotoxicity.

The impact of MR antagonism on CIS nephrotoxicity is controversial: spironolactone has been reported to exacerbate CIS-induced nephrotoxicity in rabbits (33), whereas eplerenone treatment had a protective effect in balb/c mice (34). Moreover, daily spironolactone administration for one week reduced CIS-induced nephrotoxicity caused by a single CIS injection in Wistar albino rats (35). Here, we show that spironolactone, but not eplerenone, exacerbates nephrotoxicity in the C57Bl/6 mouse during acute and chronic treatment with CIS. Eplerenone neither worsened or lessened this effect of CIS treatment. The various effects observed in our study *versus* those previously reported may be related to the timing of MR antagonist administration, as well as differences in species susceptibility but it may also relate to off target effect of spironolactone, independent of MR blockade.

In fact, it is difficult to understand how two members of the same pharmacological class can display such different effects. We show in the present study that spironolactone potentiates the deleterious effect of CIS on the cellular viability of human kidney cells in culture. This suggests that the *in vivo* potentiation of CIS is not related to a functional deleterious effect of spironolactone, such as dehydration induced by the diuretic effect of the drug, but rather a direct potentiating effect of spironolactone on CIS toxicity in epithelial cells. A major difference between the two MRAs is that spironolactone also acts as an androgen receptor antagonist or progesterone receptor agonist while eplerenone does not since it is much more specific for MR. While we cannot discard this possibility, previous results rather favor an off-target effect of spironolactone that synergizes with the cytotoxic effects of CIS. Indeed a recent study identified spironolactone in an unbiased screen of molecules which sensitized CIS-dependent cellular toxicity (24). It is well known that CIS affects various DNA-associated process, such as replication and transcription, which leads to cell apoptosis. Nucleotide excision repair (NER) is a mechanism that removes DNA lesions caused by UV irradiation and platinum-based drugs, such as CIS and is expected that the inhibition of NER would be a powerful tool to sensitize cancer cells to chemotherapy (24). Surprisingly the unbiased screening for CIS potentiating agents underlined that spironolactone indeed potentiated the effect of CIS on DNA damage through the inhibition of NER, whereas the more specific MR antagonist eplerenone had no effect (24). The authors reported that spironolactone, but not eplerenone, is a nontoxic NER inhibitor that can increase the sensitivity of CIS and UV irradiation, leading to lower viability of cells (24). Similarly, spironolactone potentiated the cytotoxicity of CIS in human pancreatic cancer cells and human colon cancer cells (36). Indeed, we translated this concept to human kidney cells, showing decreased cell viability upon CIS co-treatment.

The results of our study may have a potential clinical impact that needs to be further explored; the deleterious potentiating effect of spironolactone should be considered in cases of combined treatment with CIS and spironolactone (for example to treat heart failure or resistant hypertension), whereas eplerenone or another class of MR antagonists may be used more safely. The treatment regimen used in the chronic setting in mice in the present study was appropriate to test the prevention of repeated acute renal injury induced by each dose of CIS but do not to test the impact of repeated administration of CIS on top of long-term MRA administration as it could occur in patients. This is a limitation of our study and the impact of different treatment regimen should also be explored in a dedicated animal model.

**Table 1.**
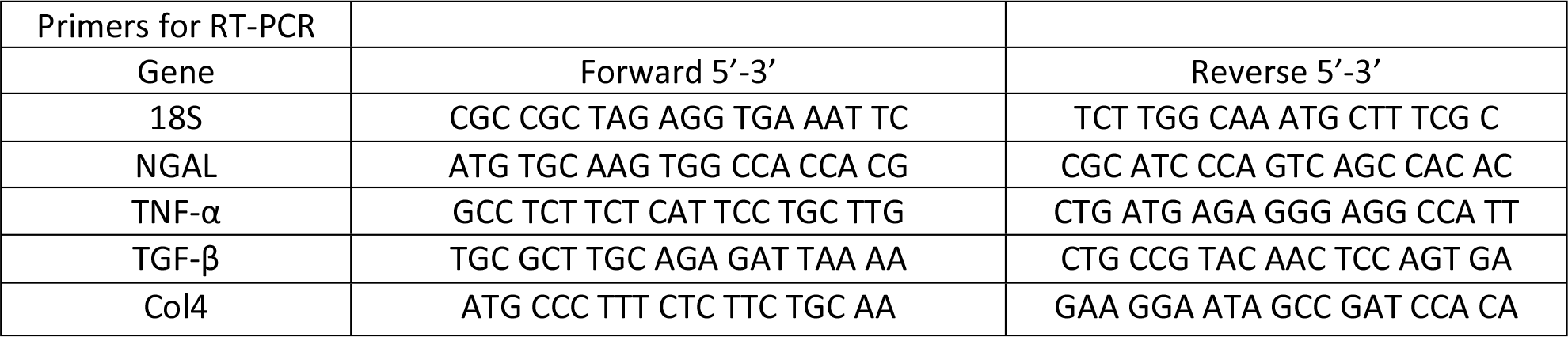
Sequences of the primers used for real-time PCR assays.

## Acknowledgements

This work was supported by grants from the Institut National de la Santé et de la Recherche Médicale, the Agence Nationale de la Recherche (ANR-16-CE14-0021-01), the French Medical Research Foundation (DEQ20160334885), the National Autonomous University of Mexico – DGAPA - PAPIIT (IA200117 and IN202919 to JBC), and the EU COST Action BM1301 European Cooperation in Science and Technology ADMIRE -Aldosterone and Mineralocorticoid Receptor (MR) Physiology and Pathophysiology (ADMIRE http://www.admirecosteu.com). BB is supported by a PhD ARDoC grant from the Region of Ile de France.

## Authorship statement

Designed the study: GRE and FJ

Performed the experiments: GRE, JB-C, and BB

Analyzed the Data: GRE, JB-C, and BB

Wrote the paper: GRE, JB-C, FJ

